# Validation of 3D cryoEM single particle reconstruction correctness and handedness with Ewald’s sphere correction

**DOI:** 10.1101/2024.08.29.610390

**Authors:** R. Bromberg, Y. Guo, D. Borek, Z. Otwinowski

## Abstract

The correct description of quantum scattering places the observed scattering contributions on the Ewald’s sphere and its Friedel mate copy. In electron microscopy, due to the large radius of the Ewald’s sphere, these scattering contributions are typically merged during data analysis. We present an approach that separates and factorizes those contributions into real and imaginary components of the image. When an inverted solution is calculated, the map derived from the real component of the image generates an inverted solution, while the map derived from the imaginary component of the image generates an inverted and sign-flipped solution. Therefore, the sign of correlation between reconstructions derived from the real and imaginary components provides the automatic determination of handedness and additional validation for the quality of 3D reconstructions. The factorization and its implementation are robust enough to be routinely used in single-particle reconstructions, even at resolutions below the limit where the curvature of the Ewald’s sphere affects the overall signal-to-noise ratio.

## 1. Introduction

The conservation of energy in elastic scattering is geometrically represented by the Ewald’s sphere ^1^. Consequently, in cryogenic electron microscopy single particle reconstruction (cryoEM SPR), the Fourier transform of the phase contrast image represents the data on the Ewald’s sphere and its Friedel mate, which is a second, inverted copy of the Ewald’s sphere, representing the complex conjugate of the wave function also contributing to measurement. Due to the very short wavelength of electrons, the Ewald’s sphere is almost flat at the resolutions of interest, and so contributions from data projected on the Ewald’s sphere and its Friedel mate can be merged on the plane that is tangential to both spheres (Figure 1A dashed line, Z = 0, Z is a reciprocal space coordinate with respect to beam direction, not to be confused with defocus *z*). For this reason, 3D reconstruction performed in reciprocal space starts from only considering information merged on the plane (the planar approximation) (Figure 1A, yellow dots) ^2-4^. However, more precise description of the projected data using a curved Ewald’s sphere can improve 3D reconstruction (Figure 1A, green and blue dots). Apart from the occasional improvements in resolution ^5-11^, we have identified additional benefits of considering Ewald’s sphere curvature, which we discuss here.

**FIG. 1.**
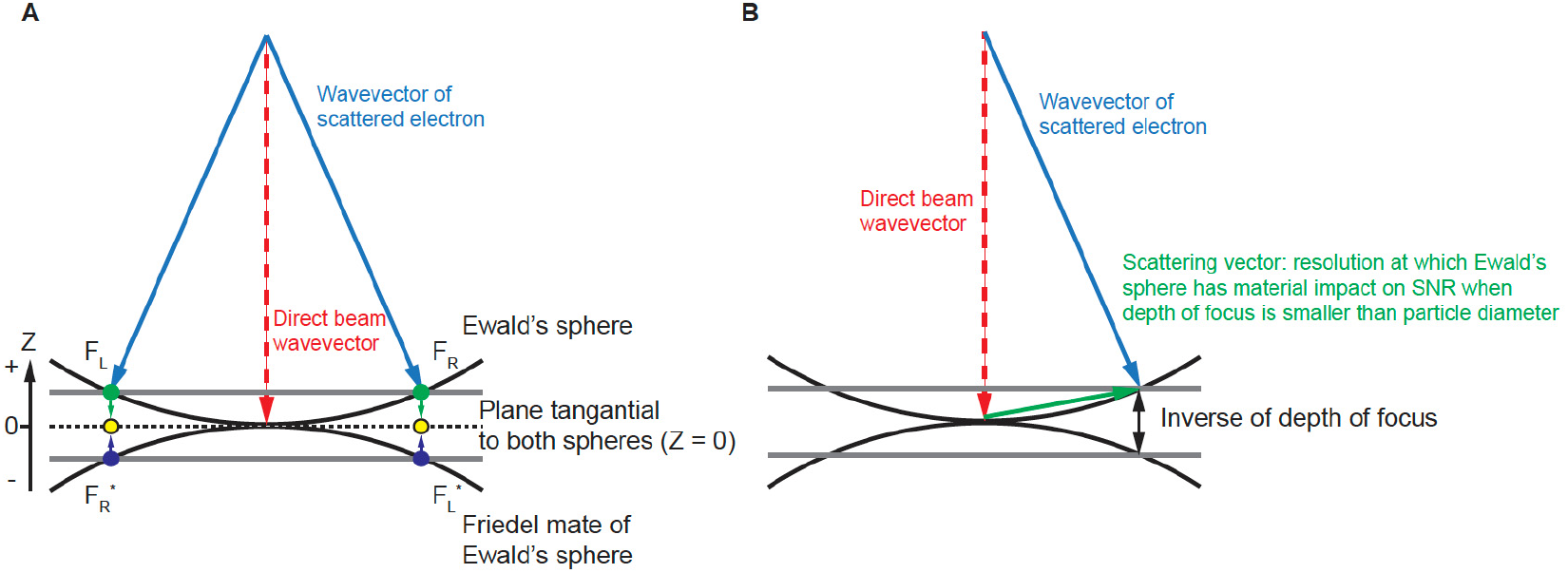
Geometry of diffraction measurement. A) Ewald’s sphere and its Friedel mate. Friedel symmetry in 3D (inversion around the origin) creates the Friedel mate of the Ewald’s sphere. We project the Fourier transform of the phase-corrected 2D image onto the Ewald’s sphere, and the complex conjugate of this Fourier transform to the Friedel mate of the Ewald’s sphere. B) The Ewald’s sphere is defined by wavevectors of scattered electrons and the separation between the Ewald’s sphere and its Friedel mate defines the inverse of the depth of focus where it has material impact on the SNR.

In a cryoEM SPR experiment, the major source of contrast in a transmission electron microscopy (TEM) image comes from interference between direct and elastically scattered electrons. A scattered electron has a complex value wave function, ***F***_***e***_, while ***F***_***d***_ represents the wave function of the unscattered direct beam (Figure 1B). By the quantum mechanics Born postulate, the measured phase contrast signal (intensity) is proportional to:

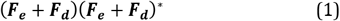

where (***F***_***e***_ + ***F***_***d***_)^*^ is the complex conjugate of (***F***_***e***_ + ***F***_***d***_). In the planar approximation that ignores Ewald’s sphere curvature, the observed phase contrast signal (aberration-corrected image in real space) results only from the real part of the wave function ***F***_***e***_ in real space, while the contribution from the imaginary component of the aberration-corrected image cancels out in reconstruction space. When Ewald’s sphere curvature is considered, both the real and imaginary parts of the wave function ***F***_***e***_ contribute to the reconstruction.

Including Ewald’s sphere curvature in cryoEM SPR improves the signal-to-noise ratio (SNR). The magnitude of improvement depends on the ratio of the particle size (particle diameter) to the depth of focus at a given resolution (Figure 1B). This ratio increases rapidly with the resolution of scattering. Depth of focus L is given by:

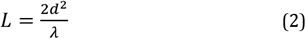

where *d* is resolution in Å, and *λ* is electron wavelength ^4, 11, 12^. When the depth of focus is smaller than particle diameter, Ewald’s sphere curvature should be considered in reconstruction. Conversely, if at the limiting resolution the depth of focus is larger than particle diameter, consideration of Ewald’s sphere curvature adds little to the SNR and its proxy, the Fourier Shell Correlation (FSC) between half-maps (half-maps FSC).

However, considering Ewald sphere curvature also provides other benefits besides increasing the SNR. It was postulated that the imaginary part of the wave function ***F***_***e***_ can provide handedness differentiation ^10^. Wolf *et al* ^10^ in the section 2.5 of their publication proposed to determine handedness by calculating correlation coefficients between observed particle images and particle images calculated from a reference structure and its inverse. However, the proposal was not implemented and doubts were raised recently ^13^ on whether handedness determination with this approach is practical. This pessimistic assessment might arise from the fact that the handedness determining signal would be low for correlations calculated for individual images of particles. We created a sensitive handedness Handedness with Ewald’s sphere correction determination method by signal factorization at the level of full 3D reconstruction and show that automatic handedness determination that considers the imaginary component of the wave function ***F***_***e***_ can be used for a broad range of projects.

We also show that this method can be used for validation of 3D reconstruction independently of other statistical indicators such as half-maps FSC ^14-16^.

### A. Algebra of reconstruction based on the Ewald’s sphere

Data acquisition happens on 2D detectors, and the first level of description of reconstruction is in the detector (2D) space. The consideration of Ewald’s sphere curvature requires expanding the description to three dimensions even at the level of a single image.

We start with the exit wave function ***F***_***e***_ = *e*^*iφ*^ using *φ* = 2π***W***, where ***W*** is the wavefront function. In the first approximation that ignores higher-order aberrations, the wavefront function is:

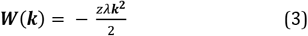

where *z* is the defocus, and ***k*** is the scattering vector. We apply our CTF based estimate of the wavefront function as a phase shift to the observed image to generate the phase shifted image ^17^. The wavefront function can also be split into base (***z***) and correction (Δ***z***), where Δ***z*** represent the defocus difference in the correction to the wavefront function.

In reciprocal (momentum) space, the exit wave function has both Hermite and anti-Hermite components which by the Fourier transform are equivalent to the real and imaginary parts in real (image) space. Experimentally, we measure interference between scattered and direct beams. This measurement contains a mixture of signals from real and imaginary components, because scattering in two opposing directions (Figure 1A, **F**_**R**_ and **F**_**L**_) creates the same periodicity ^11^. Scattering waves in opposite directions (**F**_**R**_ and **F**_**L**_) are far apart in the Fourier transform of the image (Fig. 1A, **F**_**R**_ and **F**_L_ represented by green dots). When mapped on the Ewald’s sphere and its Friedel mate, they become close (Figure 1A, **F**_**R**_ and **F**_**L**_**\***, or **F**_**L**_ and **F**_**R**_**\***, represented by green and blue dots), with their distance resulting from the contribution of the Ewald’s sphere curvature.

## 2. Methods

### A. Factorization of handedness signal

We noticed that the Hermite and anti-Hermite components of the wave function, or equivalently in real space real and imaginary parts of the image, have a different relationship to the inversion of a 3D reconstruction. The inversion of the reconstruction can be achieved by resetting particle orientation from Euler angles α (ϕ), β (θ), γ (Ψ) to α, β, γ+180°, where γ represents rotation in the image plane or equivalently the complex conjugate of the Fourier transform of the particle (Figure 2) ^18^. Note that changing α by adding constant value would rotate the reconstruction by that value, without inversion.

**FIG. 2.**
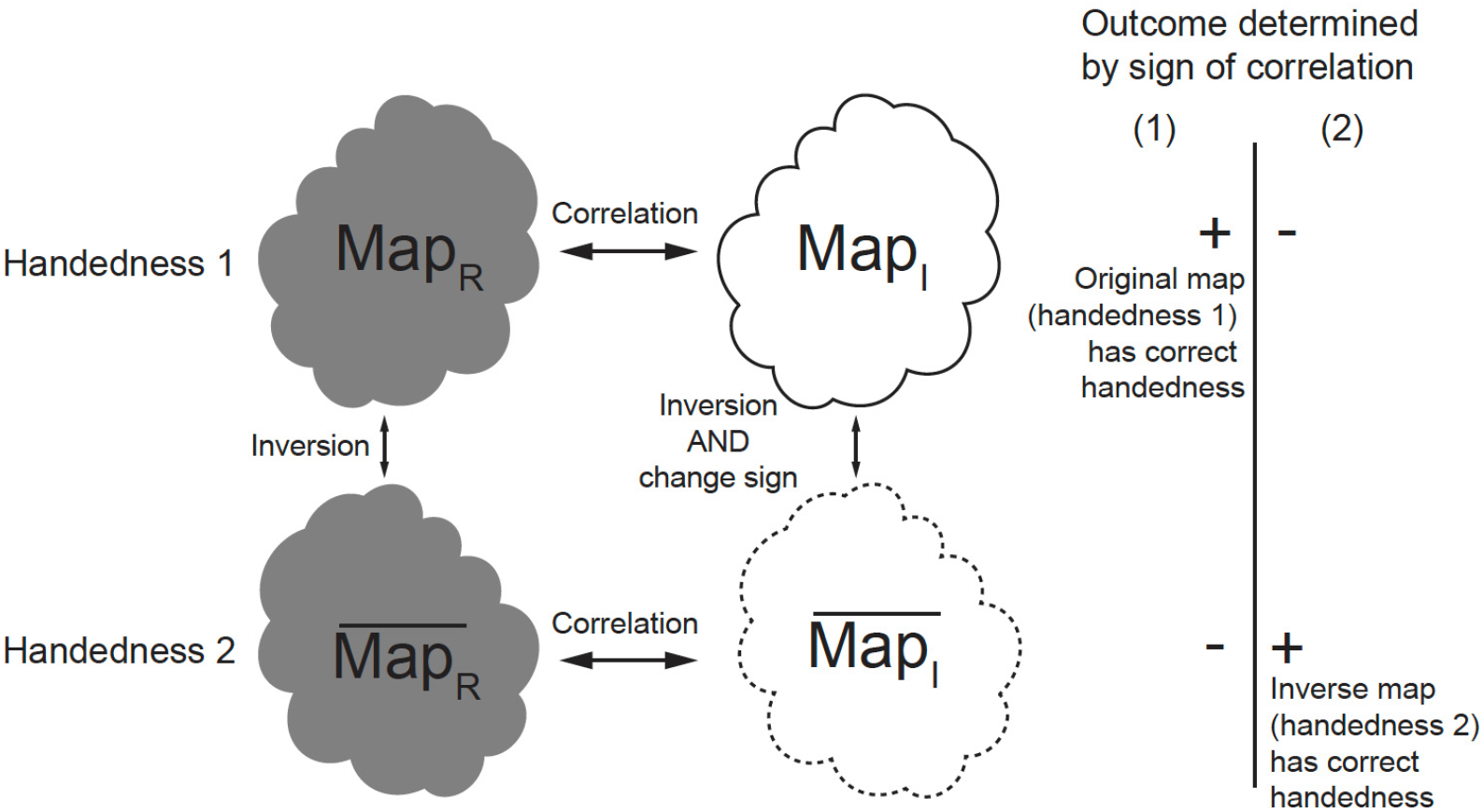
Separate reconstructions from the real component and imaginary component help handedness determination. ***Map***_***R***_ and 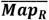 are related by inversion only. ***Map***_***I***_ and 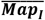 are related by inversion and flipped sign. FSC between ***Map***_***R***_ and ***Map***_***I***_ and FSC between 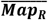 and 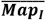 are used in handedness determination. If ***Map***_***R***_ has the correct handedness as in situation (1), the FSC between ***Map***_***R***_ and ***Map***_***I***_ will be positive; if 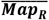 has the correct handedness as in situation (2), and FSC between 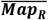 and 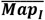 will be positive.

Map statistics (including histogram and half-maps FSC) from the reconstruction using the Hermite (real) component of the wave function (***Map***_***R***_) are identical to map statistics from the inverted reconstruction 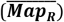. For this reason, handedness cannot be derived from inspection of ***Map***_***R***_ and 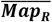 alone. However, the reconstruction from the anti-Hermite (imaginary) component of the wave function (***Map***_***I***_) changes both handedness and sign upon inversion 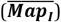 (Figure 2). This allows us to factorize the problem of handedness determination. We can separately perform reconstructions from the real component and imaginary components of images and investigate the sign of correlation between these two reconstructions (correlation between ***Map***_***R***_ and ***Map***_***I***_, and cor-relation between 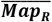 and 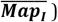 (Figure 2). The correct handedness will result in positive correlation, while the wrong handedness will lead to negative correlation, assuming the handedness of the detector has been calibrated. These correlations are informative even if the resolution is moderate because we use many particle images in reconstructions, so weak signal becomes significant when it is averaged over all of them. Typically, we have enough particles to calculate this correlation in resolution shells, so here we present it as FSC between two reconstructions obtained from the same dataset: one based on the real component and other based on the imaginary component of particle images (FSC between ***Map***_***R***_ and ***Map***_***I***_, and FSC be-tween 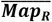 and 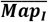). If ***Map***_***R***_ has the correct handedness, the FSC between ***Map***_***R***_ and ***Map***_***I***_ will be positive; otherwise 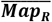 has the correct handedness, and FSC between 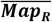 and 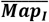 will be positive (Figure 2). When FSCs for both possibilities are plotted, the mirror feature of the FSC plot makes it very intuitive to assess if the handedness signal is significant. In principle, the highest sensitivity will be obtained by averaging correlations across resolutions, but typically even a single shell is highly statistically significant.

Because of the relationship between real and imaginary components-based reconstructions towards the 3D map and its inversion, the inverted map can be derived automatically from the original map, so we do not double the computation time.

The resulting two FSC correlation curves will be an exact mirror image of each other. It is also worth noting that here we assume a standard approach for the reconstruction so there is no particle orientation bias from using a reference that prefers one handedness over the other, i.e. the imaginary component is not used in the 3D reference at all. It is only used when the chirality of the solution is investigated.

### B. Resolution considerations

When we observe a particle at some defocus setting, we can consider the actual defocus value at each z-plane within the particle, with its +/-range being defined by particle radius. In a reconstruction that includes all possible projections, the average defocus spread can be characterized by twice the molecular radius of gyration, *R*_g_. When the wavefront change caused by z-depth variations within the particle reaches the value of ∼0.5 (**Δ*W***(***k***), defined in Eq. 3 as being in the range of +/-0.25), defocus variation starts affecting the reconstruction significantly so including Ewald’s sphere curvature becomes necessary. This happens at resolution higher than 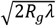. Such resolutions are achievable for some larger, rigid molecules, but are not typical.

For example, virus AAV2 has a radius of gyration of 120 Å. The resolution at which the defocus variation starts affecting resolution of the reconstruction is equal to ∼2.2 Å ^7^, assuming 300 kV electrons with wavelength of 0.019687 Å. In the case of apoferritin, which has a radius of gyration of 60 Å, the Handedness with Ewald’s sphere correction resolution at which the accurate defocus determination becomes important is 1.55 Å for 300 kV electrons. These cases are examples of where Ewald’s sphere curvature is a critical contributor during the final steps of analysis, but in the case of most other systems, we are usually well below the resolution limit where defocus variation (i.e. Ewald’s sphere curvature) is material for the result resolution.

Our goal is to achieve reliable handedness determination at resolutions lower than those discussed above. We accomplished this by factorizing the signal into its real and imaginary components and performing separate reconstructions for each component. The imaginary component contributes to the signal even at low resolution, in a way that depends on the size of the molecule. This contribution can be recognized and recovered by averaging and the more particle images are included in the reconstruction, the stronger the SNR for this contribution becomes. Since our method is applied close to the end of the reconstruction process, when the final map is calculated from hundreds of thousands particle images, the averaging effectively recovers the imaginary component, even though it is much smaller than the real one.

### C. Implementation

The numerical calculations associated with reconstructions are performed on grids in Fourier space. The grid associated with the source signal is a 2D Fourier transform of a boxed particle image corrected for CTF. The grid associated with the reconstruction is a 3D Fourier transform of two sums from 2D contributions. One sum consists of their weighted signal, and the other sum is of the weight squared. The weight originates from the product of two terms: one is associated with an interpolating kernel (Gaussian kernel in our case), the other is associated with the interference term between the Ewald’s sphere and its Friedel mate copy. The interference term corresponds to CTF in a planar approximation for real signal. The particle orientation is quite random with respect to the 3D grid, so the grid points associated with the 2D particle information fall between the grid points of the 3D reconstruction. Therefore, we need to define a method to transfer information from one grid to another. There are multiple schemes that bridge the gap between the two grids, and they all involve some type of interpolation or integration kernel, e.g., the Kaiser-Bessel Window function ^19, 20^. The reconstruction calculations are performed in reciprocal space and can start from traversing the Fourier transform of the particles (data space) and then mapping data space grid points on reconstruction space. Such methods are called scatter-based because each data space grid point can be scattered on eight or more reconstruction grid points. Alternatively, we can start from traversing the reconstruction grid point and then projecting it to data space and such methods are called gather-based ^20^ (Figure 3). Both types of methods are used, and they perform equivalently in terms of accuracy, but may have different speed of calculations depending on the implementation. We present the details of our approach using a gather-based method with a Gaussian kernel.

**FIG. 3.**
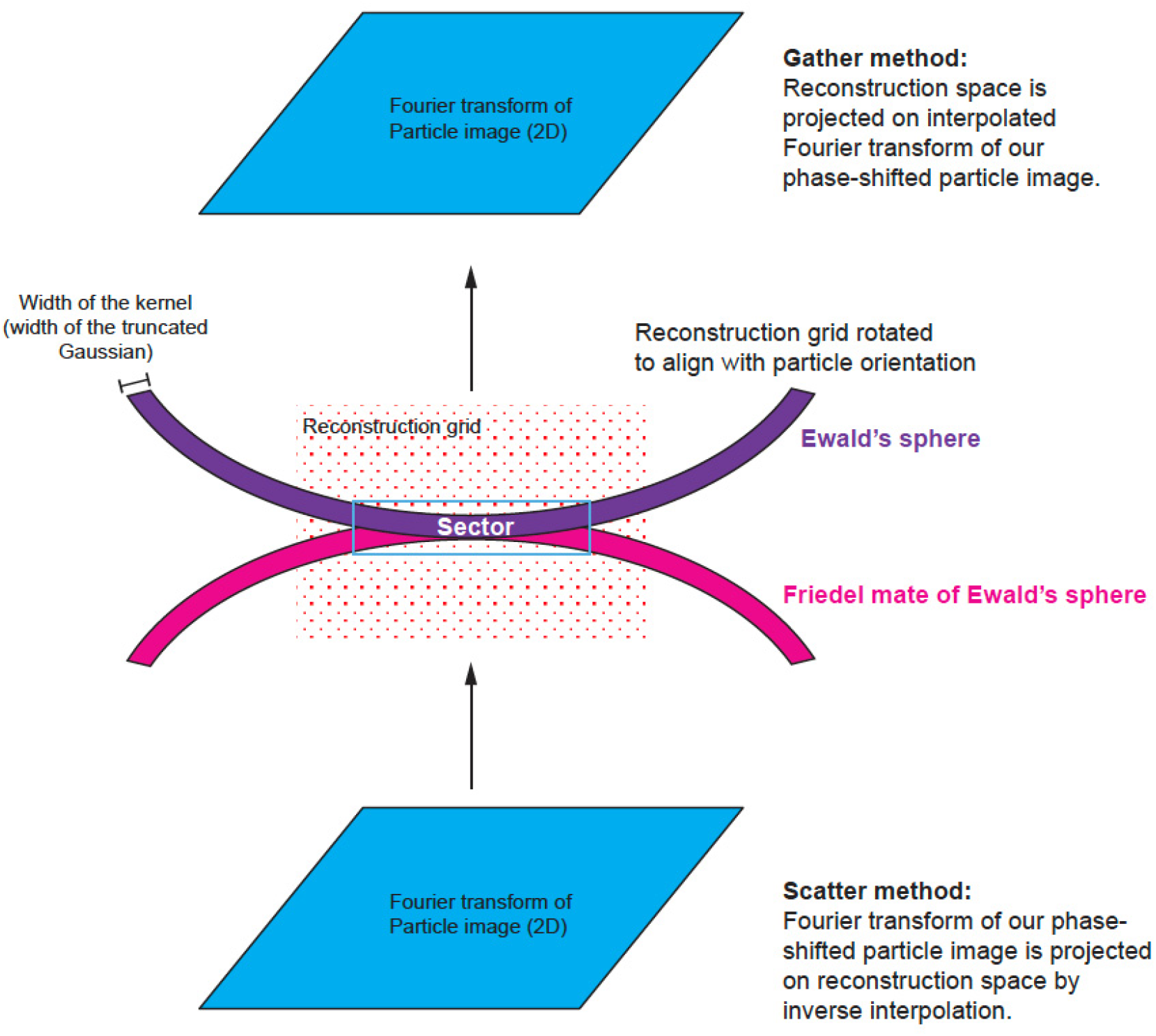
3D geometry of reconstruction from a single planar image. There are two methods that are used in the field: the scatter method and the gather method. The scatter method takes the Fourier transform and for each Fourier element, assigns the signal and its weights to the nearest points on the 3D reconstruction grid with linear interpolation weights. However, the calculation is not linear interpolation but rather the inverse of it. This is why the method is called ‘scatter’. The gather method starts by mapping each point from the reconstruction grid onto the Fourier transform of the image and then calculates an interpolated value of the image and assigns it to a single 3D reconstruction grid point. With Ewald’s sphere correction, both methods have to take into account the need to follow the surface of a sphere rather than a plane, and to consider the additional weight associated with the distance of a reconstruction grid point from the Ewald’s sphere. This additional weight may follow various interpolation kernels, e.g. linear, Kaisser-Bessel, Gaussian, etc.

We extend the gather method from Strelak *et al*. where only planar approximation was considered, to include contributions from both the Ewald’s sphere and its Friedel mate copy (Eq. 1 in ^20^):

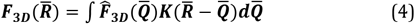

Where 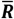 is a coordinate within a 3D reconstruction grid, 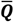 is a kernel convolution integration variable, and ***K*** is the interpolation kernel expanded to consider interference between the two spheres. 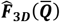 are experimental data mapped to both spheres.

We first Fourier transform each 2D particle image collected by the detector and apply CTF as a phase-shift term. We then calculate Hermite (real) and anti-Hermite (imaginary) components of the particle. Next, we accumulate separately the reconstruction signal from real and imaginary components. While the orientation of particles is the same for both reconstructions, the contribution from particle images (real vs. imaginary) and weights are very different both in the CTF term and in how the kernel contributions are summed from both spheres.

For each particle, we identify a relevant 3D sector for reconstruction that is thick enough to account for a Gaussian kernel and Ewald’s sphere curvature (Figure 3 and Figure 4). We map each point of the sector onto our particle’s data space.

**FIG. 4.**
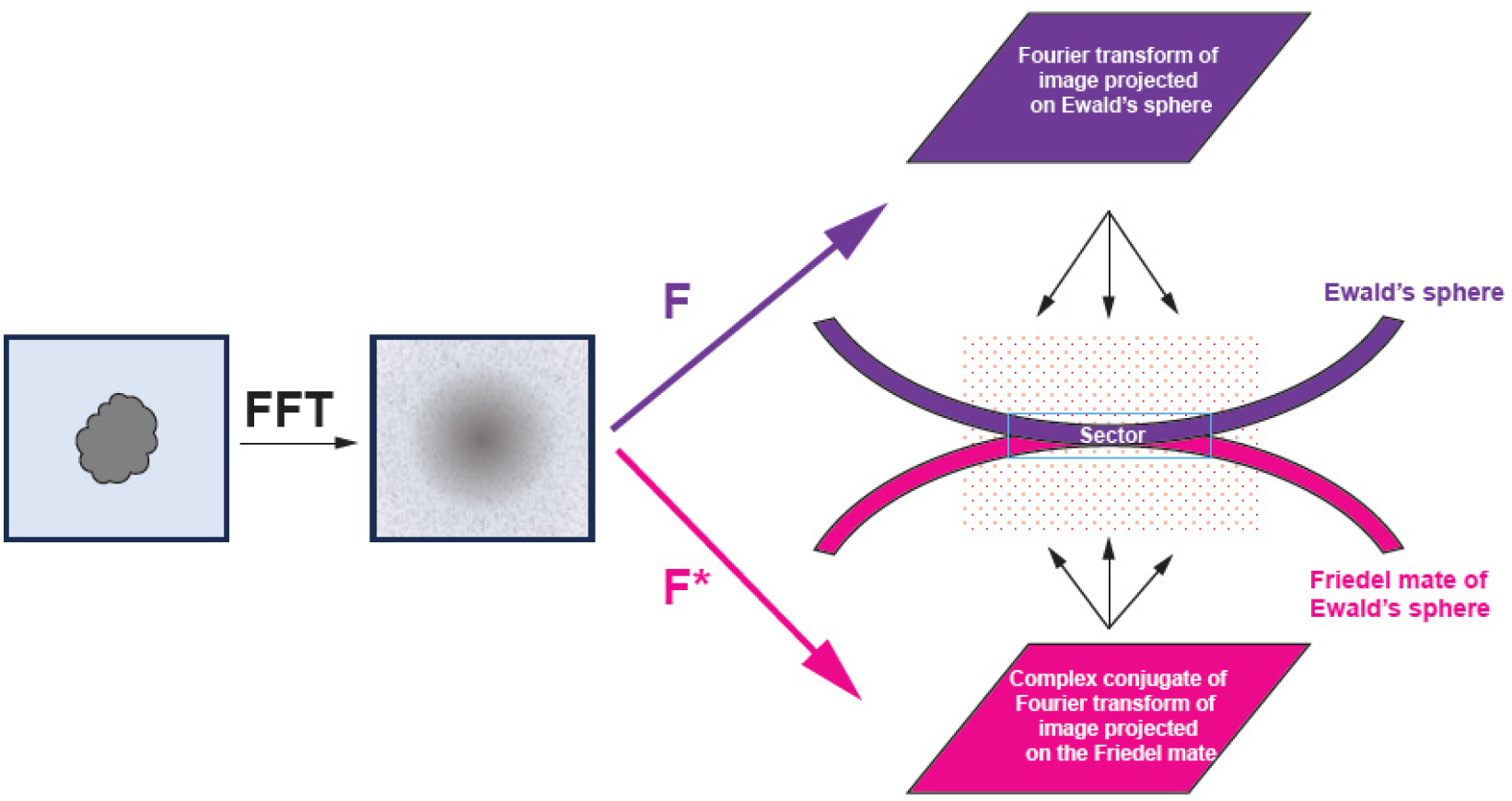
Geometry of 3D reconstruction. From an image, we calculate the Fourier transform and the complex conjugate of it. The Fourier transform (F) is mapped on the Ewald’s sphere while the complex conjugate (F*) is mapped on the Friedel mate of the Ewald’s sphere. This calculation is performed on the reconstruction grid where both spheres may contribute to the same grid point and these contributions add as complex structure factors. When these two spheres are almost touching, this addition generates a modulation known as Thon rings. Under this condition, only the real component of the image contributes to the reconstruction. Note the orientation of the 3D reconstruction grid relative to Ewald’s spheres is different for each particle.

In the gather method, the kernel defines the weight contribution from the Fourier transform of the phase-shifted detector image mapped onto the Ewald’s sphere. The weight factor is the sum of squares of the summed kernel contributions from the Ewald’s sphere and its Friedel mate.

For the real component, the kernel contributions from the Ewald’s sphere and its Friedel mate copy are always positive regardless of their distance in Z (Figure 5A, right panel (1)-(3), each kernel is represented by a dashed curve), so they contribute to the weight factor in an additive way (Figure 5A, right panel (1)-(3) solid curve with green area). The SNR is calculated from these sums by an established method ^21-24^.

**FIG. 5.**
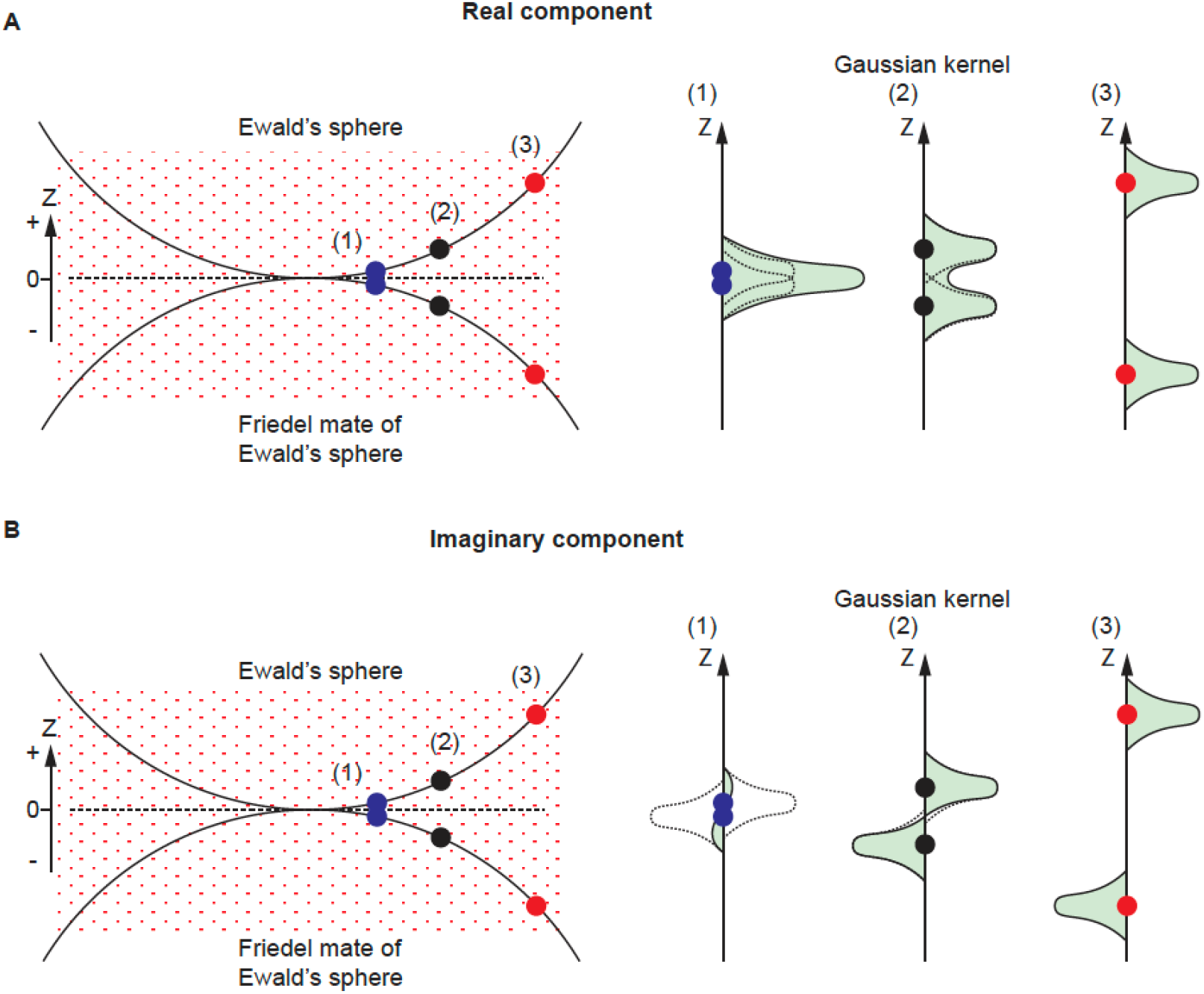
Gaussian kernel contributions for real and imaginary component. A) For the real component, the kernel contributions from the Ewald’s sphere and its Friedel mate are always positive regardless of their distance in Z (right panel (1)-(3), each kernel is represented by a dashed curve), so they contribute to the weight factor in an additive way (solid curve with green area). B) For the imaginary component, the kernel contributions from the two spheres have the same shape (Gaussian) but of opposite signs (right panel (1)-(3) dashed curve). When the reconstruction space point is close to Z=0 such as in (1) (blue dots), the two kernels will cancel each other out so that the sum of the kernel contribution at such a point is close to 0. As a result, the contribution to the weight factor (square of the summed kernel contribution) at such a point is small. For points away from Z=0, as in (2) and (3) (blue and red dots), each sphere’s kernel contribution stops overlapping with the other and the sum of the weight squared increases.

For the imaginary component, the kernel contributions from the two spheres have the same shape (Gaussian) but of opposite signs (Figure 5B, right panel (1)-(3) dashed curve). When the reconstruction space point is close to Z = 0 such as in Figure 5B (1), the two kernels will cancel each other out so that the sum of the kernel contribution at such a point is close to 0. As a result, the contribution to the weight factor (square of the summed kernel contribution) at a such point is small. This is the reason why we do not have imaginary signal contribution when using the planar approximation for reconstruction (e.g., for low resolution reconstruction). However, the sum of weight squared increases when the points move away from Z = 0, where each sphere’s kernel contribution stops overlapping with the other (Figure 5B, (2) and (3)). Similar to the treatment of the real component, the sum of the weight factor squared is the source of our SNR calculation for the imaginary component reconstruction. For this reason, the SNR for the imaginary component reconstruction is low at low resolution and improves when resolution increases, until the noise is too high.

The property of reconstruction kernels is associated with real and imaginary reconstructions. The kernels have opposing symmetry along the Z axis (Fig. 5). The kernel associated with the real reconstruction has mirror symmetry around Z=0, while the kernel associated with the imaginary reconstruction has anti-mirror symmetry around Z=0 (plane XY). The signal contribution from a single image (one particle) will have no expected correlation between its real and imaginary components as the integral of the product of any symmetric and antisymmetric functions is 0. The same applies to a group of particles that represent the same projection (only rotate in the XY plane, which is equivalent to preferred orientation) where the expected Handedness with Ewald’s sphere correction correlation between real and imaginary reconstruction is 0.

Once particles of different orientations are included in the reconstruction, the kernels of real and imaginary reconstruction associated with different particles will no longer be orthogonal, so the expected correlation between real and imaginary reconstructions will no longer be 0. The noise in individual particle images is uncorrelated, and the noise pattern of a single particle image does not introduce correlation between the real and imaginary components. Thus, there is no bias in the calculation of the correlation between real and imaginary reconstructions.

### D. Weighting of the correlation between real and imaginary reconstructions

In calculations of the correlation between reconstructions from real and imaginary components, we include an additional factor to consider how the particle size and depth of focus affect the reconstruction. The reconstruction from the imaginary component uses only out-of-focus signal (when the reconstruction point is in-focus, i.e. close to Z=0, the imaginary component vanishes). For typical resolution when the particle diameter is smaller than the depth of focus, the single particle reconstruction from the imaginary component is modulated by (**Δ*z***)^**2**^, where **Δ*z*** is the coordinate difference between a slice of the particle to the center of the particle in the beam direction in real space (same as the defocus difference for the correction to the wavefront function at that slice). As a result, assuming isotropic distribution of particle orientations, the overall reconstruction based on the imaginary component (***Map***_***I***_) will be modulated by ***R***^**2**^, where ***R*** is the distance between a point in the 3D reconstruction map and the center of this map. The reconstruction from the real component does not have this modulation, and so to make the two reconstructions more similar, we applied the ***R***^**2**^ modulation to the result from the reconstruction based on the real component (***Map***_***R***_). The ***R***^**2**^ scaling is invariant with respect to the inversion of the reconstruction 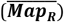, so it cannot change the sign of correlation between two reconstructions. The FSC curve representing the correlation between the reconstruction from the imaginary component and the ***R***^**2**^-modulated real component is the main result of our method. We refer to it as I-R FSC.

All data contributing to I-R FSC have been filtered with a Wiener filter^25^. Real-space unfiltered, I–R FSC curves, generated using mtriage^26^, are included in the Supplementary Materials for comparison.

### E. Consideration of kernel width

The starting point for the expected signal (SNR) from the imaginary component can be derived from either real or reciprocal space considerations. In real space, we can integrate the signal that appears on each pair of +***z*** and −***z*** planes (e.g., planes on the opposite side of the center plane of the particle in the beam direction) that increases initially with **Δ*z***^**2**^. In reciprocal space, we can calculate the second moment of the difference between two Gaussian kernels (one from the Ewald’s sphere and the other from its Friedel mate), but we have to consider that for this purpose, we should use a Gaussian kernel width that corresponds to the particle’s moment of inertia, e.g., for a small moment of inertia, we have small kernel width in real space that corresponds to wide kernel width in reciprocal space. However, in a typical reconstruction, which also applies to our real component only reconstruction, we use a narrower Gaussian kernel in reciprocal space to minimize artifacts of multiplication by Fourier transform of this kernel in real space. To simplify implementation, we also use a narrower kernel for SNR calculation and adjust the result by a compensating factor which also accounts for other departures from our over-simplified SNR theory and assumptions about the particles’ shapes, sizes and orientations discussed in the above session.

For presentation purposes, we convert the predicted SNR calculated for the real and imaginary components into a predicted FSC between the real and imaginary component reconstructions and call it predicted I-R FSC ^21^.

### F. Expected magnitude of imaginary component signal

Reconstructions from the real and imaginary components originate from the same object and therefore are inherently correlated. However, the magnitude of the correlation can vary between experiments. If we could make a prediction for the resolution-dependence of this correlation, then we would obtain a validation criterion for the correctness of our reconstruction by comparing the observed and the predicted correlations.

We use the I-R FSC and the predicted I-R FSC for that purpose. The predicted R SNR is derived from the half-maps FSC ^21^, while the predicted I SNR differs from the R SNR because different kernel weights contribute to calculations of these two quantities.

We successfully applied this validation approach, even at moderate resolution limits between 2.5 and 4 Å. We also tested a negative control using a dataset that, based on the half-maps FSC, appeared promising, but failed to produce a meaningful reconstruction. In this case, the I-R FSC showed variable signs and low amplitudes consistent with noise, and it did not correlate well with the predicted I-R FSC (see Results section for details).

This type of validation is inherently qualitative rather than quantitative, as the amount of structural disorder may depend on the distance from the center of the molecule and this disorder would attenuate the expected correlation. Also, preferred orientation would perturb this correlation by perturbing the randomness of (**Δ*z***)^**2**^ contributions to the reconstruction 3D map. The validation is best if: 1) protein is globular or spherical-shell-like, 2) particles are randomly oriented, 3) particles have no correlation between disorder and the distance from the center of the particle, and 4) the defocus error is random.

## 3. Results

### A. Handedness determination features

For each structure determination, the core result is an FSC plot between real-component-based and imaginary-component-based reconstructions (I-R FSC). For well-ordered, large molecules, the FSC can reach the level of 0.9 in some resolution shells (Figure 6A, blue curve). Interestingly, the highest correlation is reached not at the highest resolution where the signal ratio between imaginary and real reconstruction has the largest relative magnitude, but rather at intermediate resolution (∼5 Å) where the measurement accuracy is high, which likely compensates for the moderate signal ratio between imaginary and real reconstructions. The significant FSC continues down to at least 10 Å resolution. In other projects, the FSC has reached various values, with the maximum being typically at 4-6 Å resolution (Figure 6BC). The absolute configuration was unambiguously determined even if the limiting resolution was around 4 Å. We assume the method to perform well for any standard reconstruction procedure that is based on particle orientation (i.e. pose) refined against a 3D map which ignores Ewald’s sphere curvature, or alternatively using reconstruction based on the real component of the image. Under such conditions, we do not expect any model bias in the imaginary-based reconstruction.

**FIG. 6.**
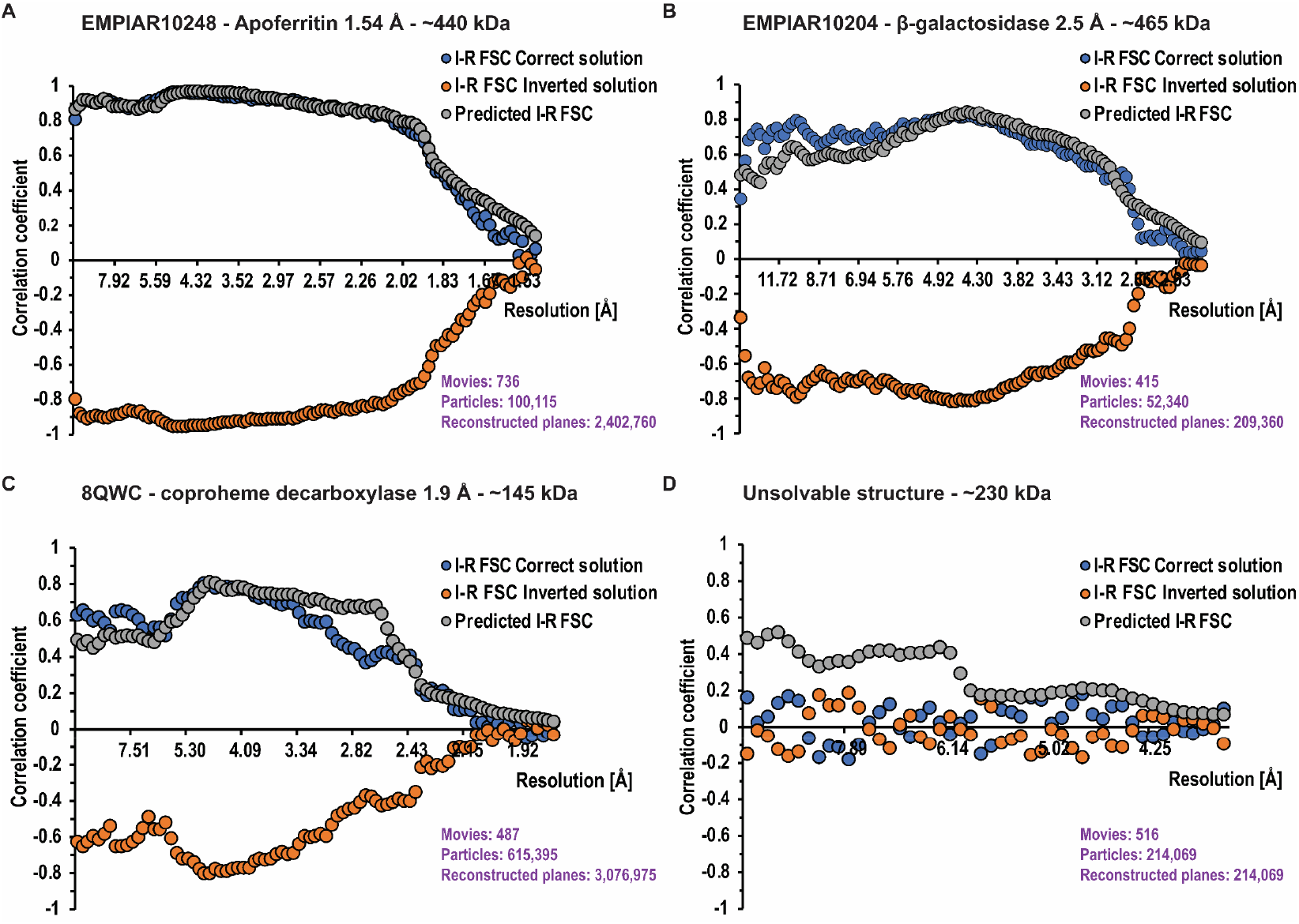
FSC statistics from the four examples. The blue and orange curves represent the observed correlations between reconstructions calculates from the imaginary and real signals (I-R FSC). There are two curves which represent the correct solution (blue, I-R FSC Correct solution) and inverted solution (orange, I-R FSC Inverted solution). The factorization of the problem results in these two curves being mirror images of each other. Separation between these curves indicates the significance of the handedness determination problem. For panels A-C, even a single resolution shell would provide more than sufficient statistical signal to indicate which solution is correct. The grey curve represents idealized predictions for the correlation when the 3D solution is right (Predicted I-R FSC). A validation criterion for the correctness of the reconstruction can be obtained by comparing the I-R FSC (blue) and the predicted I-R FSC (grey). This prediction may help to interpret the lack of separation between the blue and orange curves, to be attributed to the incorrectness of the solution (panel D) rather than the lack of statistical signal.

### B. Validation based on expected vs observed signal

For validation purposes, we investigated the relationship of the I-R FSC curves of the correct solution (Figure 6, blue curve) to the predicted I-R FSC (Figure 6, grey curve). We do not expect perfect agreement between those curves because molecules may not be globular, may have disorder correlated with distance from the molecule center, defocus determination during reconstruction may have systematic errors, and there may be preferred orientation present, among other factors affecting this agreement. Overall, we found good qualitative agreement, indicating that in the datasets we analysed, the signal from the imaginary component is properly quantified and free of major artifacts. This is important because the SNR calculated from the imaginary component is generally lower and its noise dominates the I–R FSC. We also examined imaginary component maps (I maps) in cases with strong signal. As expected, these maps exhibited stronger density at greater distances from the reconstruction center and showed weaker low-resolution features compared to the real component maps (R maps) (Figure 7).

**FIG. 7.**
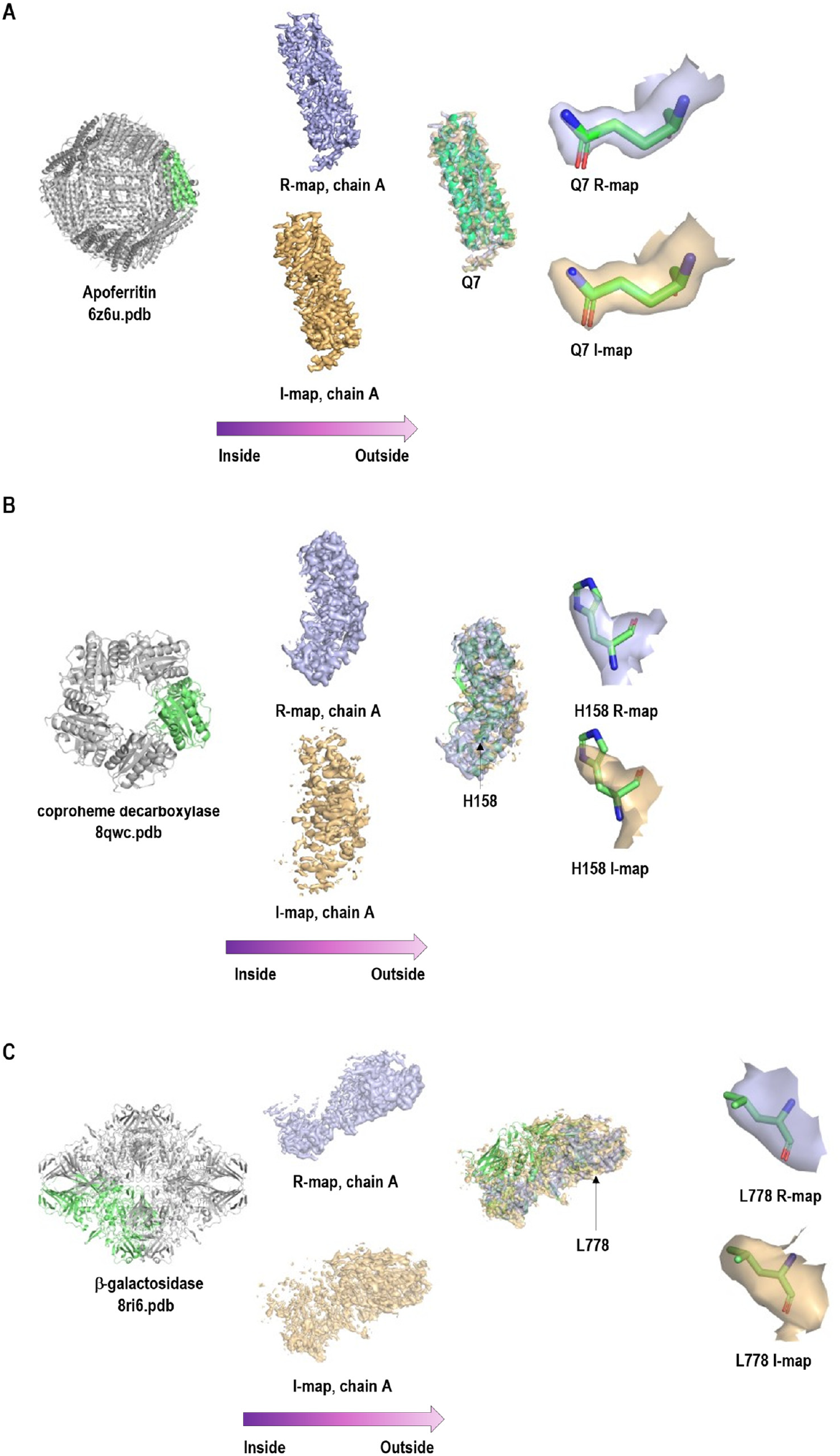
Real (R, light blue) and imaginary (I, light orange) component maps for our examples are shown. R and I maps are expected to be correlated; I maps have stronger density at greater distances from the reconstruction center and weaker low-resolution features compared to R maps. In each panel, the molecular shape is shown on the left. In the center, the magnified I (bottom) and R (top) maps for chain A are displayed, with both maps overlaid on the right to highlight density variation with distance from the center. On the far right, a magnified fragment of the I and R maps is shown to illustrate their correlation.

### C. Examples

#### Ideal case: apoferritin

We reconstructed apoferritin from EMPIAR-10248 ^27^ to 1.54 Å with O symmetry. All the core assumptions were satisfied for the case of apoferritin: the molecules are globular, preferred orientation is almost isotropic due to 24-fold symmetry, particles were of a single structural class, and the number of planes contributing to the reconstruction was very large. The I-R FSC curve went up to ∼0.9 at about 5 Å resolution and agreed very well with the predicted I-R FSC (Figure 6A).

#### Molecule of anisotropic shape: β-galactosidase

We reconstructed the β-galactosidase dataset (EMPIAR-10204) to 2.5 Å in D2 symmetry. This molecule has an anisotropic moment of inertia with an ellipsoid shape. This may explain why the predicted I-R FSC curve had some systematic mismatch with the I-R FSC depending on resolution. The resolution was about 2.5 Å with about 200,000 contributing planes, which is not a large number for a molecule with D2 symmetry (Figure 6B).

#### Small globular protein with strong preferred orientation^28^

We applied our method to a small globular protein (145 kDa) dataset reconstructed to high resolution (1.9 Å). The protein showed strong preferred orientation, and the reconstruction was done imposing a C5 symmetry. The I-R FSC curve matches the predicted I-R FSC, albeit imperfectly, most likely due to preferred orientation. The small molecule size may create some systematic errors with defocus determination per particle, which are of no significance to the real reconstruction, but may affect the imaginary signal reconstruction (Figure 6C).

#### Dataset with uninterpretable map albeit promising half-maps FSC with intermediate resolution

We also analysed a problematic case that did not converge to a proper solution in standard reconstruction, although the half-maps FSC indicated (likely overestimated) resolution of 3.8 Å (Figure 6D). The I-R FSC correlation did not show any signal, supporting the expected lack of correlation between the contribution from the real part and the contribution from the imaginary part. The expectation for a properly resolved structure (predicted I-R FSC) was above the noise level.

## 4. Discussion

We analysed the consequences of performing separate reconstructions with the real and imaginary components of the scattered (exit) electron wave. The mathematical rule is that the reconstruction from the real component has the same quality for the solution (***Map***_***R***_) and the inverted solution 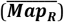, i.e. these two solutions are exactly related by inversion. However, for the reconstruction from the imaginary component, the solution (***Map***_***I***_) and the inverted solution 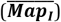 are not only inverted but also have opposite signs. The sign of the correlation between the reconstructions from real and imaginary components of particle images unambiguously determines whether the solution or its inverse is correct. This correlation is strong even for typical cases in cryoEM SPR when the reconstruction resolution is limited to ∼3-4 Å, or when particles are not too large (diameter <100 Å). Additionally, this correlation provides a new type of validation criterion. Since typical refinement schemes do not use the imaginary component of the scattered wave, there is no refinement bias towards this component, and the correlation between the imaginary component-based reconstruction and the real component-based reconstruction can only arise if the solution is correct.

Considering Ewald’s sphere curvature makes calculations more exact, but also slower and more complex. Using the traditional planar approximation is reasonable, particularly for the earlier steps of reconstruction, even for high resolution or large particles. The determination of the I–R FSC presented here could serve as the final step in the workflows for medium-to low-resolution reconstructions, followed by map inversion if the I–R FSC indicates that the handedness of the original map is incorrect. For high resolution structures, the appropriate inversion needs to be performed first, followed by reconstruction with Ewald’s sphere curvature calculations. It is important to use well-characterized handedness of micrograph images, as handedness-violating transformations (e.g., flips along the axis) can happen e.g., during conversion of image formats. Other than for this requirement, the method is reliable and does not need additional validation of handedness.

In terms of validation, the proposed method is not a replacement for a half-maps FSC. However, it is interesting from the theoretical perspective that validation can be achieved in addition to the traditional half-maps FSC. Our method would provide much stronger signal for data collected at low energy, for example with a 100 kV electron beam, where analysis following Ewald’s sphere curvature would be indicative of the reconstruction correctness at a much lower resolution. This method may also have practical utility as an indicator of whether a project is on the right or wrong track. There are data discussed in the Handedness with Ewald’s sphere correction literature ^29^ where such validation (I-R FSC) would provide a definitive answer.

## ACKNOWLEDGMENTS

Author contributions were as follows. R. Bromberg, Y. Guo,D.Borek and Z. Otwinowski developed the approach presented here; R. Bromberg and Z. Otwinowski implemented the approach; D. Borek acquired data; R. Bromberg, Y. Guo, D. Borek and Z. Otwinowski analysed the data; R. Bromberg, Y. Guo, D. Borek and Z. Otwinowski wrote the manuscript.

The work presented here was partially supported by National Institutes of Health (NIH), The National Institute of General Medical Sciences (NIGMS) (grant No. R43GM148105 to RB; grant No. R44GM137671 to RB; grant No. R35GM145365 to ZO); The U.S. Department of Energy (DOE), The Office of Science (grant No. DE-SC0021600 to RB); The National Institutes of Health, The National Institute of Allergy and Infectious Diseases (contract No. 75N93022C00035 to DB). The Cryo-Electron Microscopy Facility (CEMF) at UT Southwestern Medical Center has been supported by grants RP220582 from the Cancer Prevention and Research Institute of Texas (CPRIT).

This report was prepared as an account of work sponsored by an agency of the United States Government. Neither the United States Government nor any agency thereof, nor any of their employees, makes any warranty, express or implied, or assumes any legal liability or responsibility for the accuracy, completeness, or usefulness of any information, apparatus, product, or process disclosed, or represents that its use would not infringe privately owned rights. Reference herein to any specific commercial product, process, or service by trade name, trademark, manufacturer, or otherwise does not necessarily constitute or imply its endorsement, recommendation, or favouring by the United States Government or any agency thereof. The views and opinions of authors expressed herein do not necessarily state or reflect those of the United States Government or any agency thereof.

## DATA AVAILABILITY STATEMENT

The data that support the findings of this study are available within the article and its supplementary material. I and R maps for each example have been deposited in Zenodo at https://doi.org/10.5281/zenodo.14291830.

## Supplementary Figure

**FIG. S1.**
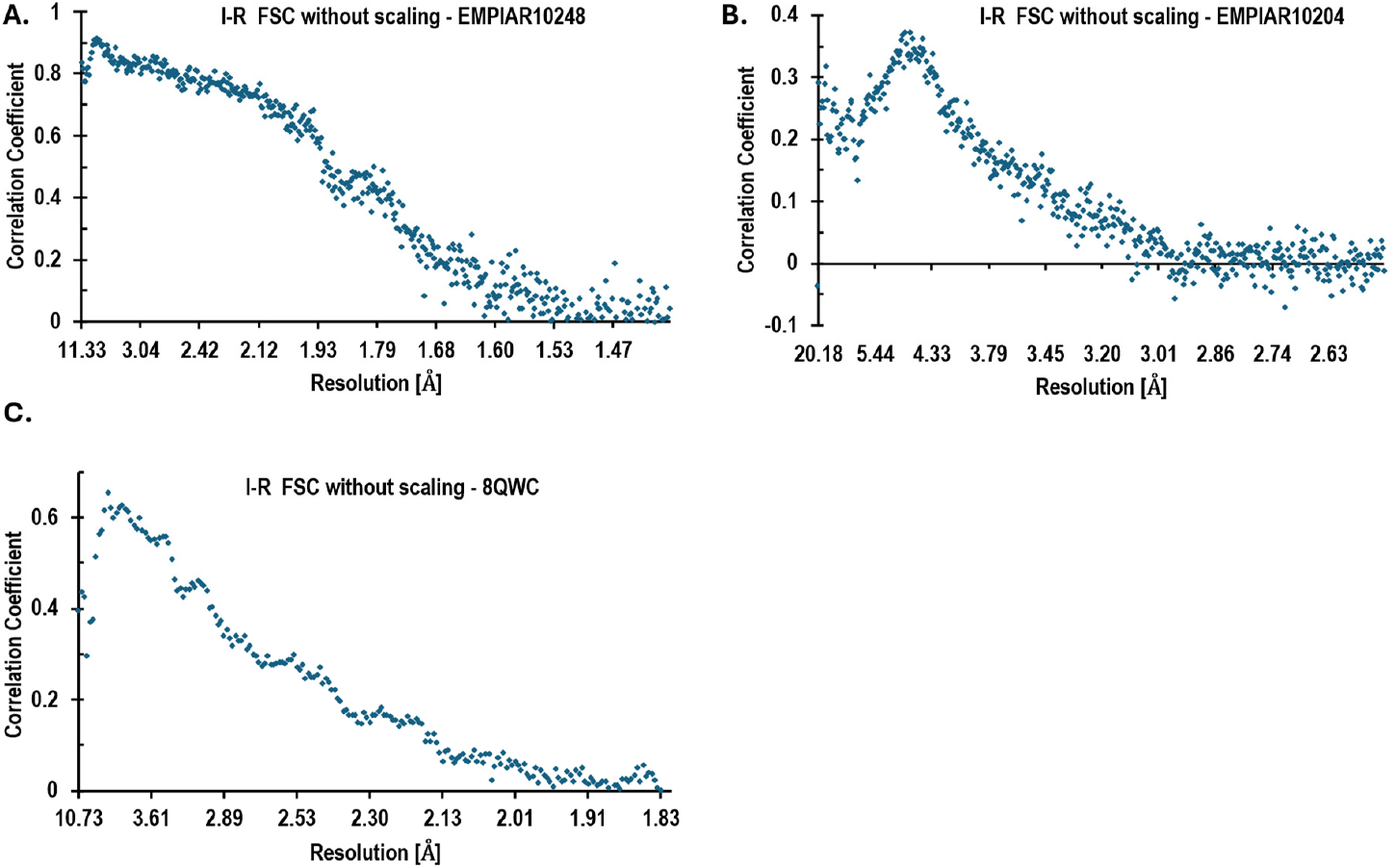
I–R FSC curves calculated on data not filtered by the Wiener filter. As expected, the correlations between I and R maps are lower for unfiltered data.

